# FishFeats: streamlined quantification of multimodal labeling at the single-cell level in 3D tissues

**DOI:** 10.1101/2025.09.02.673708

**Authors:** Gaëlle Letort, Tanya Foley, Ilona Mignerey, Laure Bally-Cuif, Nicolas Dray

## Abstract

**Summary:** Characterizing the distribution of biological marker expression at the single cell level in whole tissues requires diverse image analysis steps, such as segmentation of cells and nuclei, detection of RNA transcripts (or other staining), or their integration (e.g., assigning nuclei and RNA dots to their corresponding cell). Several software programs or algorithms have been developed for each step independently, but integrating them into a comprehensive pipeline for the quantification of individual cells from 3D imaging samples remains a significant challenge. We developed FishFeats, an open-source and flexible Napari (Sofroniew *et al*. 2025) plugin, to perform all of these steps together within the same framework, taking advantage of available and efficient software applications. The primary core of our pipeline is to propose a user-friendly tool for users who do not have a computational background. FishFeats streamlines extracting quantitative information from multimodal 3D fluorescent microscopy images (smFISH expression in individual cells, immunohistochemical staining, cell morphologies, cell classification) to a unified “cell-by-cell” table for downstream analysis, without requiring any coding. Our second focus is to propose and ease manual correction of each step, as measurement accuracy can be very sensitive to small errors in the automatic process.

*Availability and implementation:* FishFeats is open source under the BSD-3 license, freely available on github: https://github.com/gletort/FishFeats. FishFeats is developed in python, as a Napari plugin for the user interface. Documentation is available in the github pages: https://gletort.github.io/FishFeats/.

## 1 Introduction

Recently, tissue biology has become single-cell, and tremendous efforts have been placed on developing experimental set-ups to link cellular transcriptomics with cell morphology, spatial position within a tissue, and other cellular features, all at the single cell level. Many techniques in molecular biology have been developed to allow for multiplexed labelling on cells, embryos, or tissues, often in situ, for RNA transcripts at the single molecule level (smFISH) together with immunohistochemistry (IHC) staining for membrane, nuclear, or cytoplasmic components. However, extracting quantitative information from the acquired microscopy 3D images involves several image analysis steps. The number and efficiency of available tools to perform each of these image analysis steps are increasing but with it also the complexity of installation and usage of these tools (Schlaeppi *et al*. 2022; Li *et al*. 2023; Nogare *et al*. 2023; Pylvänäinen, Jacquemet and Marcotti 2025). This makes it difficult for users who do not have a computational background to take advantage of these tools and perform their own analysis, especially if it requires combining them.

We present here our open-source and user-friendly pipeline, FishFeats (for Fluorescent In Situ Hybridization Features), to facilitate the joint analysis of multiple quantitative cellular features (mRNA transcripts, IHC markers, morphometric measures) in 3D at the single cell level in epithelia (Figure 1A). FishFeats proposes a single interface to run several state-of-the-art algorithms as well as its own specific algorithms. Importantly, all layers of quantification are unified into a single table ‘per-cell’, suitable for downstream analysis. We strongly focused on its user friendliness through its graphical interface (Figure 1B) and dedicated documentation, which are important aspects for the tool accessibility (Jamali *et al*. 2021; Pylvänäinen, Jacquemet and Marcotti 2025). The frequent feedback from its users was fundamental to identify and correct usability bottlenecks (Pylvänäinen, Jacquemet and Marcotti 2025; Soltwedel and Haase). Particularly, one crucial component in all our biological applications was the possibility for manual correction. This is a tedious task that we now assist with different visualization options, dedicated shortcuts, or automated functions.

**Figure 1.**
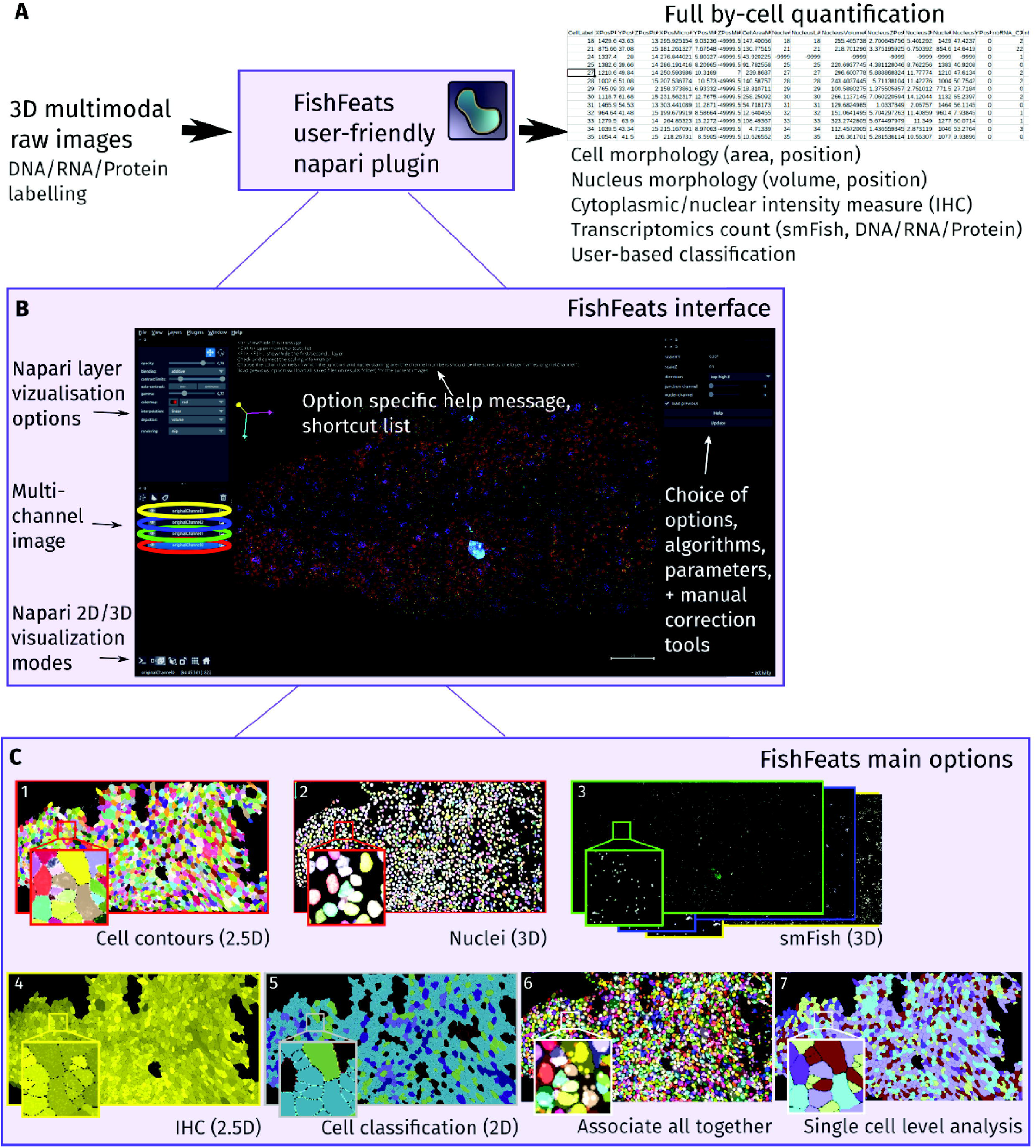
**A –** FishFeats workflow. FishFeats inputs are raw 3D multimodal images. The main output consists of a single cell-by-cell table with all the measured features combined. **B –** FishFeats main user interface, relying on Napari interface. FishFeats options are available on the right part of the interface, with all choices for algorithms and parameters. A specific help message for each option is overlaid in the main window, stating the main actions/shortcuts for this step. **C –** Examples of the main options. Snapshots of the results of each of the main FishFeats image analysis steps: 1: cell segmentation, 2: nuclear segmentation, 3: smFISH segmentation, 4: cytoplasmic measurement (IHC), 5: cell classification, 6: cell-nucleus-spot assignment (“associate all together”), 7: hierarchical analysis (“single-cell analysis”). For each panel, analysis dimensionality is provided in parentheses. 2.5D refers to 3D analysis that are not fully 3D in their implementation. The biological image showcased in this Figure is described in Experimental Data in (Supp. 4).

## 2 FishFeats

We chose to implement our pipeline as a Napari plugin (Sofroniew *et al*. 2025) which is open-source, Python based, compatible with most image formats, and importantly for our purpose, has multiple visualization advantages (2D/3D, multilayered) allowing for more interactivity. Several steps are proposed (Supp. 2) as independently as possible for flexibility in the analysis. Raw images and their metadata with standard format can be directly imported in the plugin. In most analyses, the first step consists in segmenting cells within a tissue. In developmental biology, cells are routinely identified by an apical marker of cellular junctions forming sort of 2D sheet within a 3D image (e.g. IHC for Zona Occludens 1, ZO1). FishFeats first does a projection and 2D segmentation of the cell contours and then places these labels in 3D (Figure 1C-1). The projection can be calculated within the plugin or by an external software (LocalZProjector, CARE) (Weigert *et al*. 2018; Herbert *et al*. 2021). FishFeats offers an interface to perform the segmentation of these epithelium-like projections with Cellpose and Epyseg (Aigouy *et al*. 2020; Stringer *et al*. 2021). The generalization capacity of CellPose, especially the CellPose-SAM version (Pachitariu, Rariden and Stringer 2025), makes it compatible for use with a wide range of non-epithelial cells. Manual correction of the results is facilitated by Napari Label editing tools and additional specific options such as keyboard shortcuts to switch channels on/off, merge cells, etc. (Supp. 2.1.2). The 2D cells are then re-positioned in 3D by assigning to each cell its most likely Z-position, which can be fine-tuned or manually corrected (Supp. 2.2). If the acquired images contain nuclear staining (or any structured labelling, such as vesicles), 3D segmentation can be done (Supp. 2.3) using Stardist or CellPose (Weigert *et al*. 2020; Stringer *et al*. 2021) (Figure 1C-2), along with the possibility to filter and correct the results. Each segmented nucleus or object can then be paired with the apical cell surface it likely belongs to with the Hungarian algorithm (Kuhn 1955) (Supp. 2.4). Manual corrections of the cell-nucleus or cell-object association are made easy by clicking on the image with keyboard shortcuts (Supp. 2.4).

A core development of the pipeline is the feature to detect and assign single RNA molecules (visible as dots) to their corresponding cell. Segmentation of RNA dots is performed in 3D (Figure 1C-3) with Big-Fish (Imbert *et al*. 2021), a robust spot detection library. Important parameters of the algorithm can be tuned in the interface. If needed, segmented spots can be manually edited (deleted or added) thanks to Napari Point layer tools and customized shortcuts (Supp. 2.5). Assigning these dots of single RNA molecules to their corresponding cells defined by their 2D apical areas is notably complicated as cytoplasmic markers are often not different enough between neighboring cells to determine the full cell volume. Thus, we implemented several strategies to compute an automatic assignment of 3D spots to their cell. The most obvious assignment method is the Z-projection of spots to the cell apical segmentation. We also propose an assignment based on the closest nucleus, or by mixing the two approaches depending on the depth of each spot to the apical surface, by defining a convex hull of the volume of the cell or iteratively by using previous RNA assignments (see comparison of approaches in Supp. 2.5.2). In a dense tissue context, manual correction of automatic RNA assignment is necessary to measure correctly the cell genetic profile (Mitchel *et al*. 2025). However, it is also a quite tedious task. We put extra effort in proposing convenient tools for manual correction of RNA assignments based on frequent feedback from the biologists (Supp. 2.5.2).

## 3 Additional features

To reduce the number of channels required experimentally when combining IHC and/or DNA/RNA labeling, we have incorporated a tool in FishFeats that permits the computational separation of nuclear staining from an apical junction marker if both are co-stained within the same channel. FishFeats can create two virtual channels containing each biological signal, either based on morphological filters or on a home-made deep learning solution. The latter solution, called SepaNet, was implemented with a U-Net-like architecture, that we separated in two output upsampling parts after the latent space calculation (Supp. 2.8). While our neural network is specific to junction-nuclei separation, it could be easily retrained to separate other structures. Recently, other deep learning architectures have also been proposed to implement similar separations (Jin *et al*. 2024; Ashesh *et al*. 2025), which might be interesting to test on our images in future developments.

To quantitatively describe cells based on a cytoplasmic level of expression we have built a tool that, from the apical segmentation, can measure the intensity of any channel within a user-defined volume below the plane of the cell (Figure 1C-4). Importantly for many biologists, we also added the possibility to add manual annotation. These classifications can be automatically pre-filled from the cytoplasmic intensity of any channel (Supp. 2.9; Figure 1C-5) and/or manually based on the user expert eye. Here the user can click on cells to assign them identities. Finally, all these features (cell/nucleus morphologies, transcriptomics content, cytoplasmic or nuclear intensity measures, classification) are joined in a single cell-by-cell table (Figure 1A), ready for downstream analysis. We added an option in FishFeats to directly perform hierarchical clustering of the cells from that table (Figure 1C-7) (Supp. 2.9).

## 4 Conclusion

Nowadays, a lot of very powerful image analysis tools are available for a given task. However, for a non-expert user, finding the right tools, installing and using them, and combining their results together can be quite challenging. The gap between the development of the tools and the accessibility to the user is further increasing with the success of deep learning solutions. Moreover, most analyses require manual corrections, as well as the creation of ground truth datasets to train new deep learning algorithms. There is thus a strong need for more accessible tools in the bioimage analysis field (Carpenter, Kamentsky and Eliceiri 2012; Schlaeppi *et al*. 2022; Pylvänäinen, Jacquemet and Marcotti 2025; Soltwedel and Haase). We initially developed FishFeats to internally deploy a user-friendly tool for our specific multimodal spatial analyses of tissues at the single-cell level, and are now proposing this development as a flexible and accessible tool to the community. Features in FishFeats were mainly added when we encountered their need. We expect that our pipeline will continue to grow in the future and, importantly, will be maintained in the coming years.

## Experimental Data

Experimental data used to showcase our plugin in this Application Note and its supplementary data are described in the supplementary data file (Supp. 4).

## Supplementary Data

Supp.: Supplementary File “supdata_fishfeats.pdf”

## Acknowledgement

We thank the members of the Zebrafish Neurogenetics team of Institut Pasteur for their insightful feedback on the plugin, and Romain Levayer, Jean-Yves Tinevez, and all the members of the image analysis hub of Institut Pasteur for discussions on the plugin development.

