## Supplementary material for "FishFeats: streamlined quantification of multimodal labeling at the single-cell level in 3D tissues": Supp.

**3 Results table** **14**

**4 Experimental data** **14**

### 1 Presentation

#### 1.1 Installation

*FishFeats* is deployed as a pip module and can be directly installed in a python environment by typing: `pip install fishfeats`. It is implemented as a Napari [Sofroniew *et al.*, 2025] plugin to use its interface for visualization and user interactions.

See the documentation page https://gletort.github.io/FishFeats/Installation for full installation instructions and version compatibilities.

#### 1.2 Usage

To start the pipeline, launch Napari in your virtual environment where *FishFeats* is installed (`napari`), and go to `Plugins>FishFeats>Start`.

Once you have started the plugin, a file dialog opens to let you choose the image to analyse. The currently accepted formats are `.tif, .czi, and .ims`. The image will then be displayed with the different color channels shown as separate layers in Napari (left bottom interface). In the right part of the interface, the options specific to *FishFeats* are displayed. The first step is to set-up the image scaling properties and to indicate in which channel the apical junction staining (+/− nuclear staining) are. Then the plugin lets you decide which step to perform (Figure 1).

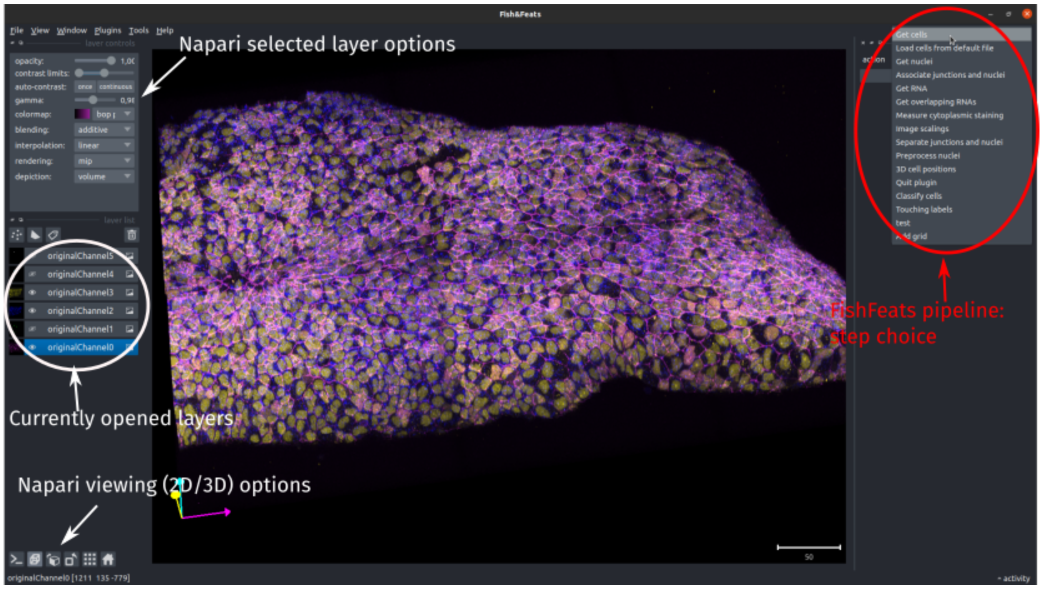

**Fig 1.** *Main interface of the FishFeats plugin, with the viewed 3D image in the center, the Napari panel on the left and the plugin options in the right panel*

For ease of use, *FishFeats* proposes several shortcuts (additionally to those already available in Napari) specifically for each step of the pipeline. At all steps, you can press `<h>` to show or hide the current step shortcut and informations. You can also refer to the step wiki pages for a more detailled description of the shortcuts and step usage.

For usage flexibility, all steps are available from the start and can be performed at any time. However, be aware that some steps require that other steps are run beforehand. An error message would then appear indicating to choose the required prior step.

*FishFeats* is evolving following user feedbacks and biological questions for which it is used. In future developments, other steps or options will be included, please refer to the repository documentation pages for more informations.

### 2 Main steps

#### 2.1 Cell segmentation

*Full online documentation here*

This step allows to determine the apical contour of the cells, based on an apical junction staining (e.g. ZO1). The segmentation is done in 2D, thus the junctional staining first has to be projected.

NB: if for experimental reasons the junction and nuclei staining were acquired in the same channel, the projection and segmentation results will be improved if the two staining are first separated. For this, FishFeats proposes an additional step to separate the two stainings (see 2.8).

##### 2.1.1 Projection

The apical junction staining can be projected in a 2D plane by a local average projection around the slice of maximum intensity of the signal within the FishFeats plugin (Fig 2). It is also possible to perform the projection with dedicated softwares such as LocalZProjector [Herbert *et al.*, 2021] or CARE [Weigert *et al.*, 2018], and load the projection directly in the plugin.

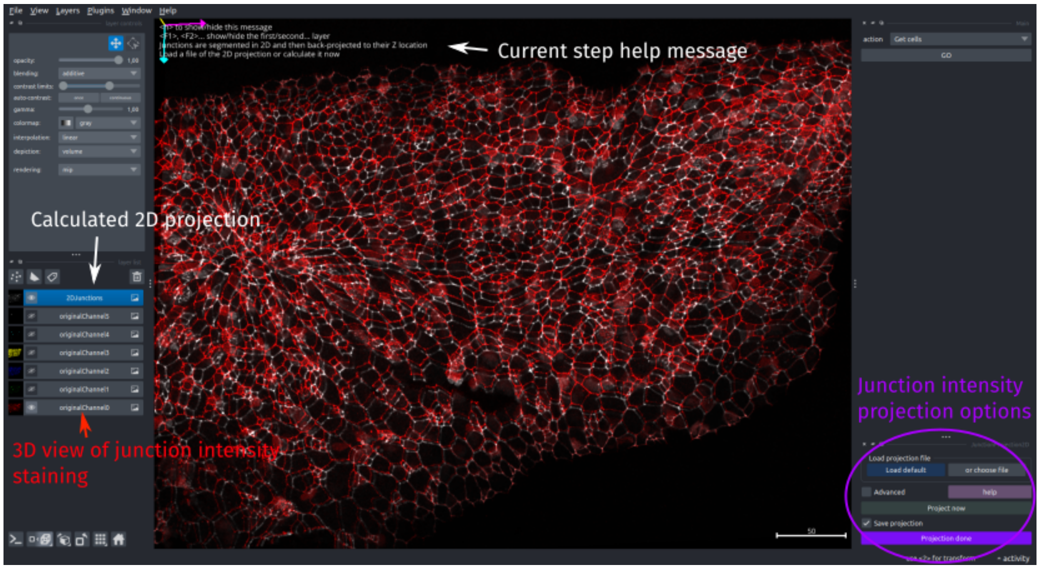

**Fig 2.** *FishFeats interface to do the 2D projection (results shown in white) of the 3D staining of apical cell junctions (shown in red).*

##### 2.1.2 Segmentation

Segmentation of epithelia can be done in FishFeats either with Epyseg [Aigouy *et al.*, 2020], or CellPose [Stringer *et al.*, 2021], or can be loaded from another source. In our experience, these two softwares are usually very efficient for epithelial segmentation and can be retrained within their framework if necessary.

Moreover, CellPose remarkable generalisation capacity, especially since CellPose-SAM version [Pachitariu *et al.*, 2025], makes it compatible also with non epithelia cells.

Nonetheless, there is still a small error rate (Fig 3). If you need a perfect (or near perfect) segmentation, FishFeats proposes several tools to ease the manual correction of the results, based on the Napari Label layer capabilities. You can paint/erase/fill/merge the cells/relabel

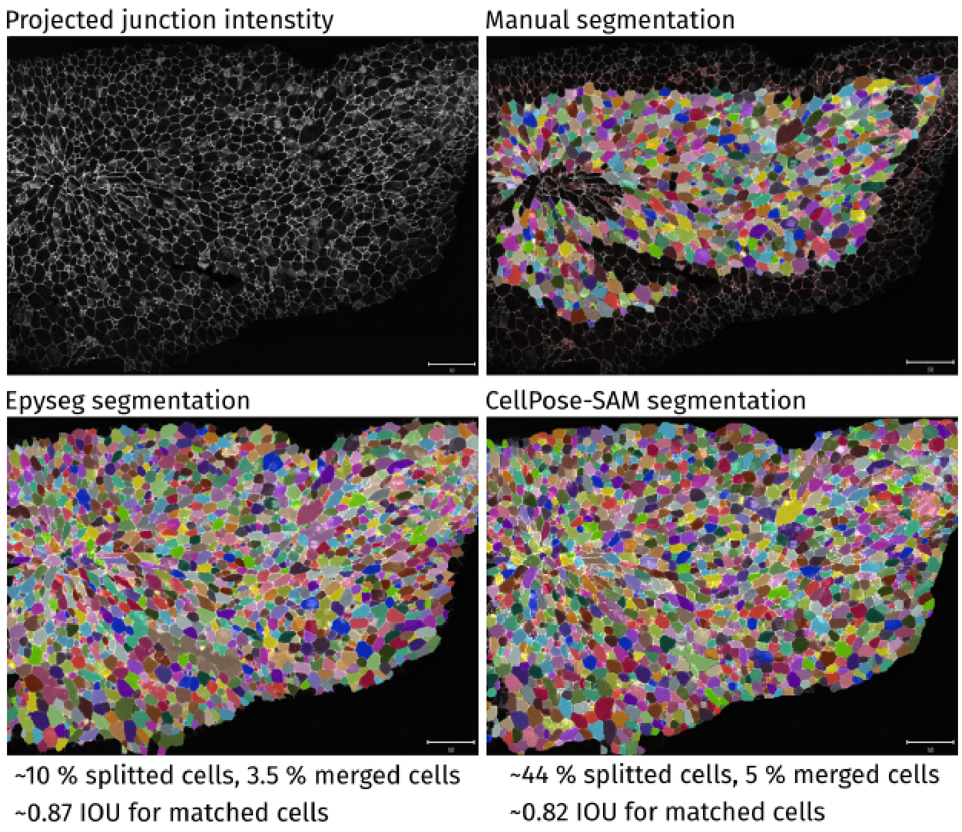

**Fig 3.** *Results of cell segmentation with the different FishFeats option (manual segmentation, Epyseg, Cellpose). The manually drawn cells were compared with the cells found in the automated segmentation to evaluate the methods performance on this image. Splitted cells=cell from the manual segmentation that was considered as several cells in the automated segmentations. Merged cells=cells from the manual segmentation that were considered in the automated as within a much bigger cell. IOU=Intersection Over Union, calculated for cells that were matched between the manual segmentation and the compared segmentation.*

the cells/change the visualisation... thanks to Napari Label tools and FishFeats specific shortcuts/options.

Here, we compared the automated segmentation results of Epyseg and Cellpose (here we used Cellpose-SAM version [Pachitariu *et al.*, 2025] but other versions can be installed instead) with manually drawn cells. The 3 segmentations were done within *FishFeats* . With both methods, the overall contour corresponded well enough with the manual segmentation (Intersection Over Union $> 0.8$) when the cell was correctly matched between the two segmentation. On this image and compared to the manual annotation, Epyseg performed much better than CellPose-SAM. Both methods had a higher tendency to split cells (i.e., consider that one cell is two or more cells, Fig 3). We thus added in *FishFeats* a `merge cells` shortcut that allows to easily merge two cells into one and thus speeds-up the manual correction of splitted cells.

#### 2.2 3D position

ℹ*Full online documentation here*

As cell segmentation is done in a 2D projection, we then have to find the position of each cell in 3D (its Z position). For this, FishFeats calculates a map of the local Z position of the

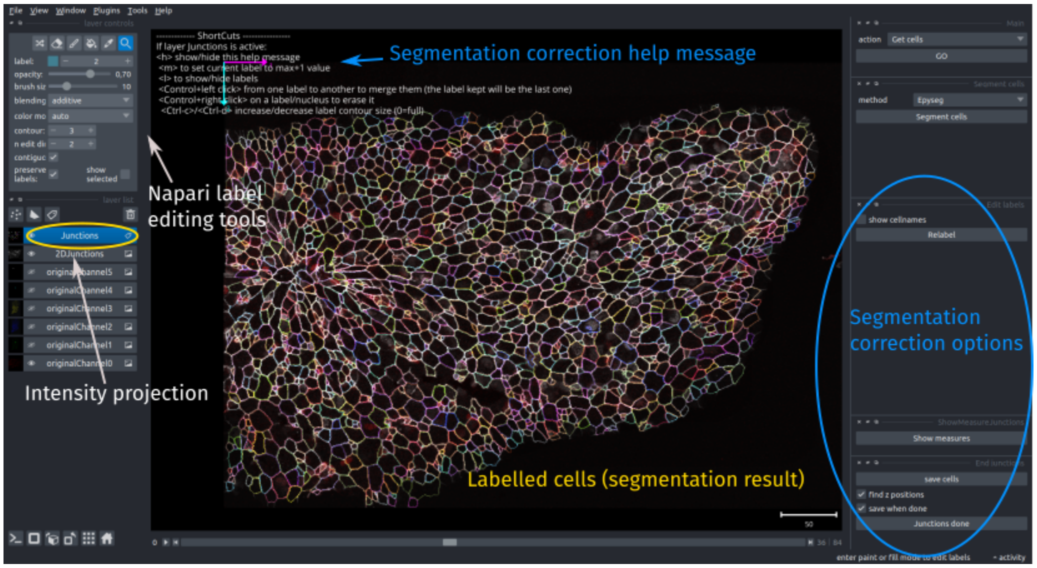

**Fig 4.** *FishFeats interface to do manual correction of cell segmentation. Segmentation results are displayed with a Label layer. Label edition tools can be used as well as some specific shortcuts from FishFeats.*

junction staining by searching for the local maximal similarity of the projected signal with the original 3D signal. This height map is constructed by sliding overlapping windows in which the projected image similarity is compared with the same window in all Z-slices. Cells are then allocated a location based on the value of the height map. Futhermore, this height map can be exported in an output file, so that it can be used in softwares that can estimate the apical cell exact 3D size (which can be miss-estimated due to the projection when the tissue has high curvature), such as DeProj [Herbert *et al.*, 2021].

The computed cell position and the precision of the height map can be changed in the FishFeats 3D position options, which give access to the main parameters of the calculation (Fig 5). Moreover, several shortcuts are proposed to facilitate the manual correction of the cells' placement in 3D (see the online documentation for more details).

#### 2.3 Nucleus segmentation

*Full online documentation here*

For nuclei 3D segmentation, *FishFeats* includes two standard options, `Stardist` and `CellPose` [Weigert *et al.*, 2020, Stringer *et al.*, 2021, Vijayan *et al.*, 2024, Kleinberg *et al.*, 2022].

`Stardist` can be used in 2D and 3D. Some trained 3D models are available, but that are less general than Stardist2D, due to the lack of publicly available annotated 3D data, the effort it demands to generate new ground-truth for the training and the model being larger by adding one dimension. Additionally, the images to analyse have different xy/z resolution anisotropies that would need to be corrected and homogeneized to perform a direct 3D analysis. Instead we decided to first focus on taking advantage of the high performance of `Stardist` in 2D and reconstruct together the nuclei in 3D by linking them as soon as they overlapped enough.

`CellPose` has two different 3D modes, one doing 2D segmentation then reconstructing in 3D (similar to our Stardist option) and a second one running segmentation on xy, yz and xz slices, then computing the nuclei shape from the calculated flows together.

If the image is large, `CellPose` computing can fail if it excesses GPU memory (e.g. more

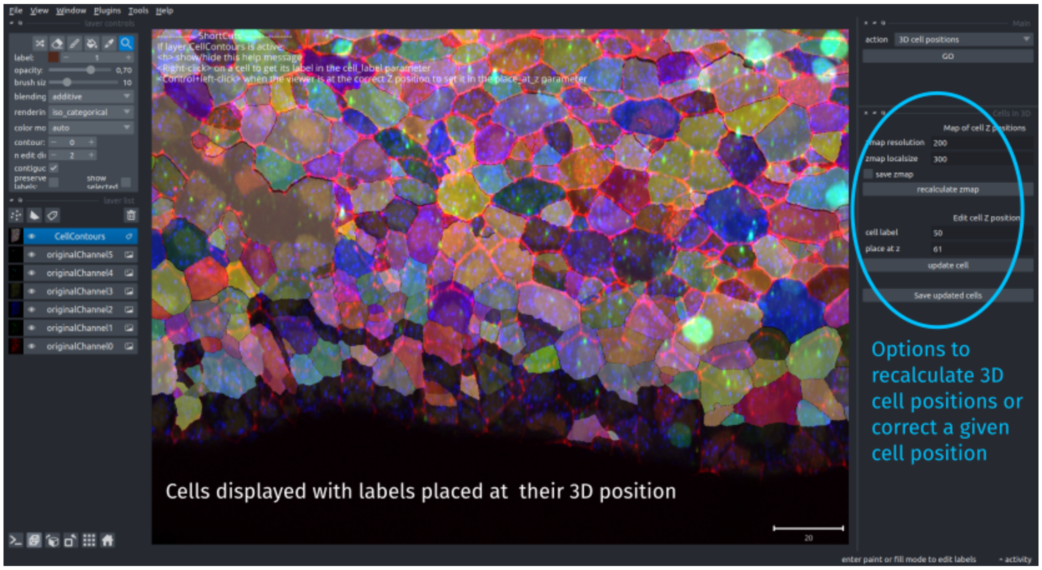

**Fig 5.** *FishFeats interface to recalculate cells 3D position or manually correct individual cell position.*

than 20 GB, depending on the GPU card). To handle big stacks, *FishFeats* has a parameter `Use dask` in `CellPose` computation that uses a custom code to run CellPose in parallel chunks with dask, inspired from the distributed segmentation contributed code in cellpose version 2

In the example shown in Figure 6, segmentation of a stack of size (85, 2871, 1950) pixels took around 15 min with Stardist2D+reconstruction, 14 min with CellPose-SAM in 2D+reconstruction mode and 60 min with CellPose-SAM in 3D mode.

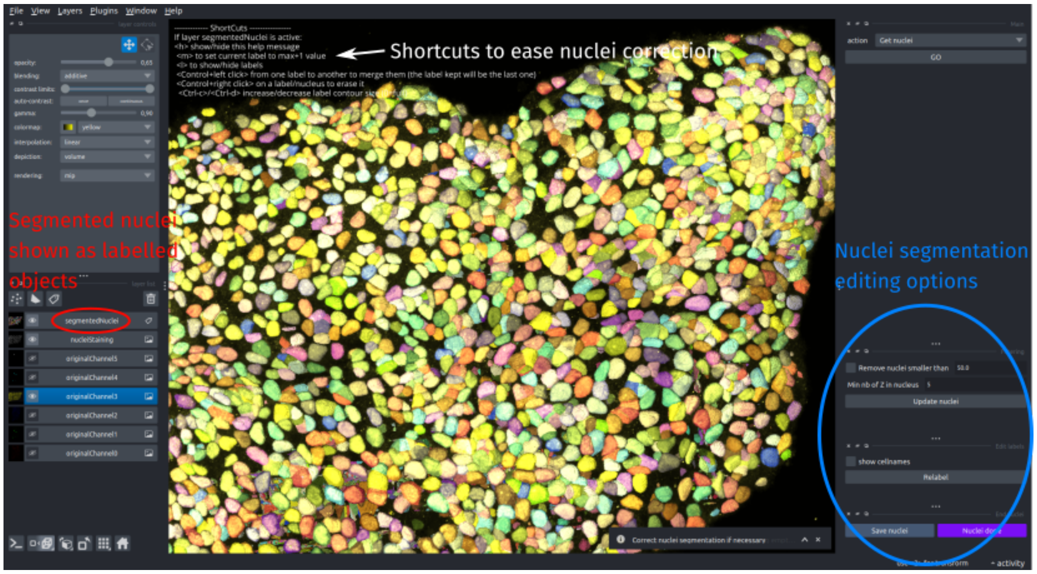

**Fig 6.** *Segmentation and correction options of nuclei in 3D. The displayed segmentation (as 3D labelled object) was calculated with CellPose in 3D mode.*

As for all the pipeline steps, some options and shortcuts are added in *FishFeats* to ease the manual correction, a painful step in general.

Nuclei can be filtered by their size to remove segmentations that are too small, merged together to correct over-splitting, or easily removed and the visualization options can be rapidly switched directly from the keyboard (Fig 6). In addition, the label edition tools of Napari Label layer are available.

Nuclei 3D segmentation and annotation are still a challenge. New algorithms and trained models are starting to be available in the community, as for example [Pop *et al.*, 2013, Wu *et al.*, 2021, Vijayan *et al.*, 2024, Achard *et al.*, 2025] for the segmentation, and [Leggio *et al.*, 2019, Borland *et al.*, 2021, Garcia-Lopez-de-Haro *et al.*, 2025, Arzt *et al.*, 2022] for manual correction. We will test the promising new solutions and integrate some in our pipeline or make it compatible with them.

#### 2.4 Cell-nucleus association

*Full online documentation here*

To reconstruct full 3D cells, nuclei should be associated to the apical cell they belong to. For this, *FishFeats* calculates an optimal pairing between the segmented cell (2D segmentation + Z position) and 3D nuclei segmentation. The pairing cost is defined by the distance between the apical cell centroid and the closest point of the nucleus surface. To find the optimal cell-nucleus pair combination, *FishFeats* uses the Hungarian algorithm [Kuhn, 1955].

After computation, nuclei are associated to the most likely cell. Their label will be set to the same label as their associated cell, so they will be displayed in the same color in the *FishFeats* interface (Figure 7). This helps to directly visualize eventual mis-association and correct it manually thanks to dedicated options (Figure 7). When the assignement of one nucleus is modified in the interface to be linked to another cell, if this cell had already another nucleus associated, this nucleus will be automatically unassigned and relabeled.

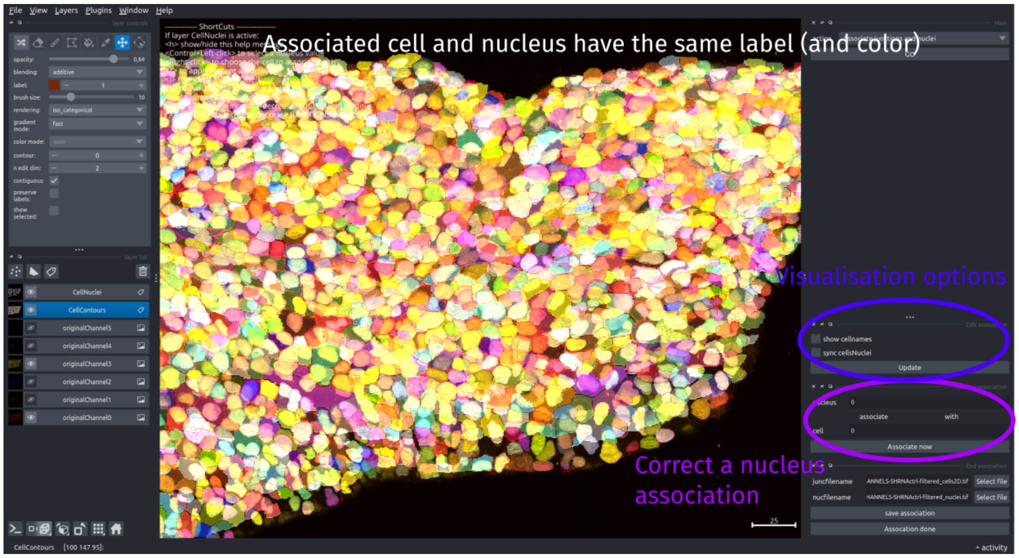

**Fig 7.** *Cell-Nucleus association step and correction options. Associated nuclei and cells are colored similarly, with only a reduced opacity of cell surface for visualization.*

Some nuclei can be not assigned to any cell if they were too far to be assigned or if there are more nuclei than cells. In that case, they have a label not assigned to any cell. Most analysis are cell-based so they will be left out of these analysis, but some steps, as e.g. measuring nuclear intensity, can still be performed on these unassigned nuclei if desired.

#### 2.5 RNA transcripts

*Full online documentation here*

##### 2.5.1 RNA segmentation

Detection of RNA transcripts is a crucial step of the pipeline, to correctly quantify the cell transcriptomic identities. Segmentation of RNA spots in 3D is done in FishFeats with Big-Fish [Mueller *et al.*, 2013, Imbert *et al.*, 2021], with which we always obtained satisfying results in our images. Big-fish is based on detecting local intensity maxima after a laplacian of gaussian filtering so a few parameters, such as the size of the spots or defining a threshold, have to be tuned (refer to the online documentation for the parameter description). Other methods based on deep learning solutions have been recently developed and have the advantage of not requiring parameter choices but might require retraining when the images are different from the training data, as for example U-Fish [Xu *et al.*, 2024]. We might integrate them in future developments of the pipeline if the segmentation step becomes more challenging or upon request.

After the spots position has been calculated, FishFeats displays them as 3D points with the point layer of Napari (Fig 8). Their displayed properties (size, visibility of some of them..) can be chosen through our interface panel and points can be added/deleted with Napari point tools.

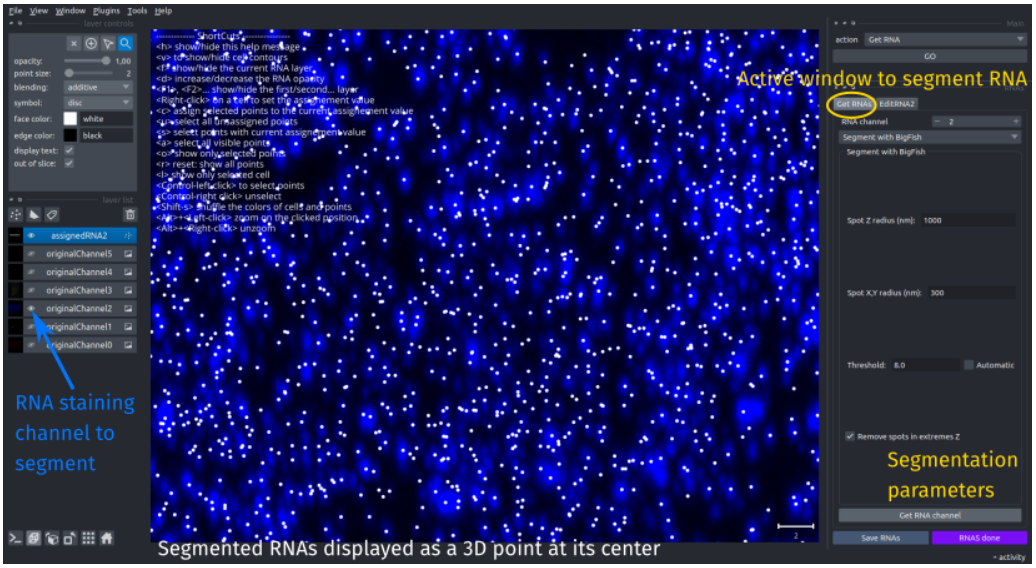

**Fig 8.** *Interface to segment transcripts for a given channel (in blue) or load already segmented spots. Detections are shown as Napari 3D points (in white).*

Once the RNA segmentation is correct, the detected points can be each assigned to the apical cell it likely belongs to.

##### 2.5.2 RNA-cell assignment

Identifying to which cell a RNA spot belongs to is not trivial in our context. Indeed, the spot are positionned in 3D. While the cell apical contour can be stained with marker as ZO1 or cadherins and thus quite robustly segmented (see 2.1), the volume of the cell is much harder to determine as we did not a marker of the cell full membrane and cytoplasmic stainings would not differ enough between neighboring cells. Thus, we propose several initial assignment methods that would have to be manually corrected when the quantification needs to be very fine.

- The most obvious assignment method is by direct projection (`Projection`) of the spot X,Y positions to the apical cell surface. However, some cells can be quite tilted compared to the apical surface and thus will be strongly misquantified by this method. Also, cell that are deliminating from epithelial tissue can have a very small apical surface while they will have a bigger volume inwards in the tissue.
- Thus, we also proposed to assign RNAs based on their proximity to cell nuclei (`ClosestNuclei`). This method is less sensible to cell area and Z-position, but necessitates a good nuclei segmentation and association to its apical cell. Moreover, RNA spots that are close to the surface will be more mis-quantified with this approach.
- Then we added a method that would mix these two approaches (`MixProjClosest` by assigning the spots close to the apical surface by projection and assigning deeper spots to the closest nuclei. This method is less sensitive to the cell position and size but requires very good cell, nuclei segmentation as well as their correct association.
- We proposed a stricter method, `Hull`, based on an approximation of the hull surface from the apical surface to the corresponding nucleus. By interpolating the surface at each Z between the apical surface and the nucleus mid-plane, all the spots inside the interpolated surfaces should belong to the cell. This method will assign less spots in general to the cells, but the assigned spots are more likely to be correctly assigned.
- Finally, an additional iterative option was implemented (`FromNClosest`). When one or more RNA channel has already been treated and assigned, we take advantage of this information and assign the next channel based on the clouds of points of the previous channel, by looking at the closest neighbors assignement. The more channels has been done before, the more reference points will be available for the new spots assignement, so the more robust it will be.

Overall, the choice and the precision of the method will depends on the specificity of the tissue and of the RNA staining (some are more ubiquitous, some are more expressed in large cells...). The projection method has the advantage of not necessiting nuclei segmentation and is quite often a good initial start. Then as RNA channels are being process, the iterative method is in general quite soon worth it. In figure 9, we show an example of comparison of the different methods, except the iterative one. In this context, the projection is very close to the manual annotation but this could be related to the expression of the tested RNA.

When the first automatic assignement has been made, if precise quantification is necessary, manual correction should be done to check and correct all spots assignement. This is a quite tedious step. We try to propose convenient options in *FishFeats* to assist in this step and makes it less painfull. An important part of this is of course to have a good initialization with the automatic methods. Then having a lot of display options to be able to easily switch the view (showing/hiding fast some layers), displaying only the cell contour, changing the size of the spots based on some properties... helped the smoothness of the process by avoiding some back and forth with the mouse pointer. The proposed options are presented in the online documentation.

#### 2.6 Cytoplasmic measurement

ℹ*Full online documentation here*

This option allows to measure the intensity (mean, median, max or min) of selected cytoplasmic stainings for each cell. As the full volume of the cell is not known but the apical surface is known, the intensity is measured only in a few slices close to the apical surface, inside the segmented apical cell. The result is a heatmap in which each cell is filled with a unique value (e.g. the mean normalised intensity, Fig 10). This option is particularly useful to measure immunohistochemical stainings and assign an identity to each cell based on the cytoplasmic

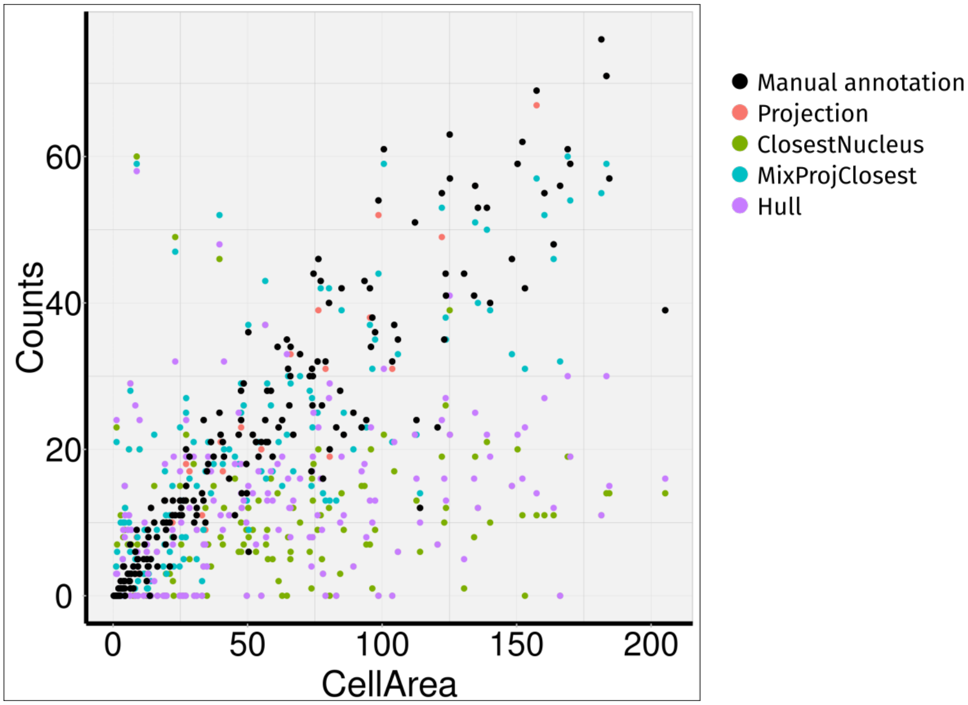

**Fig 9.** *Comparison of RNA counts in each cell with our different assignement methods relative to their apical area. Counts from a manual assignement (black) are compared to the counts from the Projection (red), ClosestNuclei(green), MixProjClosest (blue) and Hull (purple) methods. The assignements from the automated methods were not corrected. The nuclei segmentation and cell-nucleus assignement were not manually corrected, which could improved the results of some methods.*

intensity of a given marker. *FishFeats* allows to combine together the immunohistochemical stainings measurements with the cell's RNA transcript counts and thus to match cell transcriptomes with cell types.

#### 2.7 Cell classification

ℹ *Full online documentation here*

In most of our user projects, one main output was the classification of the cells into specific types/behavior/genetic profiles. Indeed, it allows to test potential correlations between the spatial arrangement of the tissue or the cell morphologies with the gene expressions or cell types.

*FishFeats* proposes an interface to easily perform this classification for different features. For flexibility, users can name their features (classification criteria, e.g. PCNA-positive cells, mitotic cells..) and include several features. The program can initiate an automatic classification based on the intensity of one selected staining or initialize a homogeneous classification and let the user determine each cell class. Indeed, the criteria for classification can be very different depending on the project, so we focus on proposing convenient editing tools.

For a given feature, the cells are displayed colored by their class. When creating the classification, the user defines how many classes are possible. Each cell can very easily be assigned to a given class thanks to shortcuts to select the current class and apply it to one cell (Fig 11).

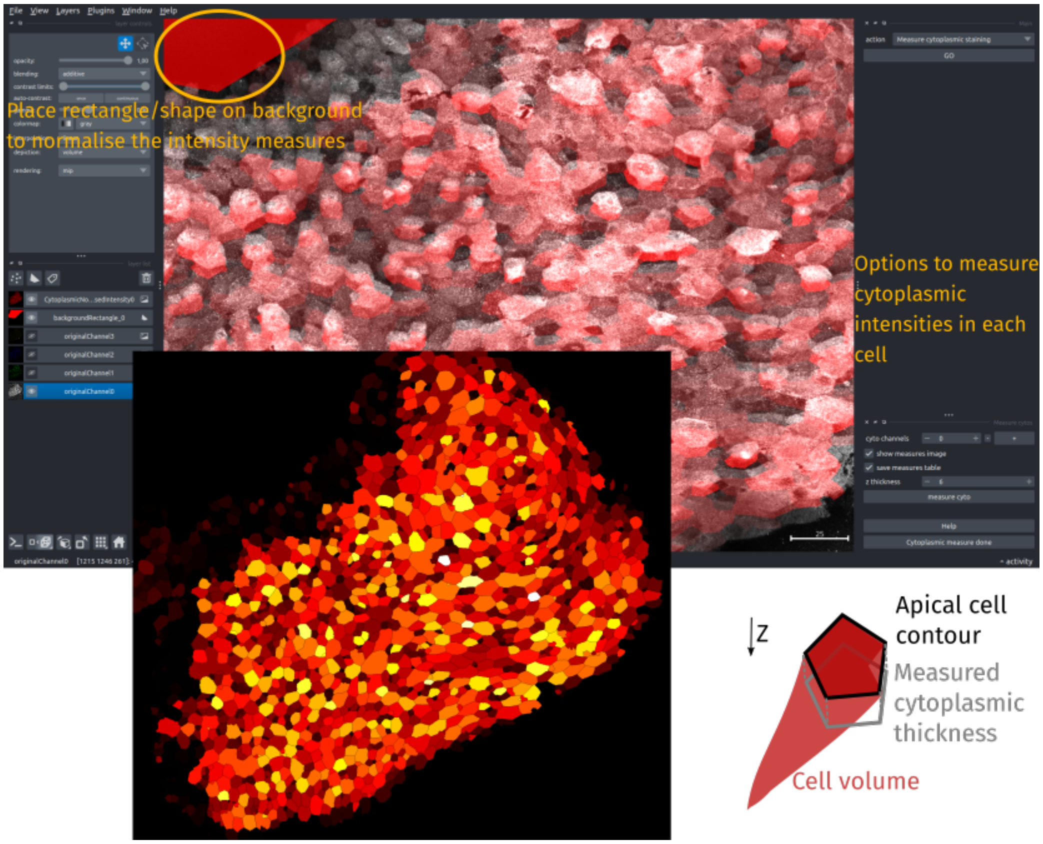

**Fig 10.** *Interface to measure cytoplasmic intensity of selected channels. The measure is done for each cell in a small volume just below the apical surface (schema in the bottom right). Rectangles (which can be deformed) allow to normalise the intensity by a background mean intensity. The results are heatmap images of each cell mean normalised intensity (bottom left image). The full table of measurement is added to the main results table.*

#### 2.8 Staining separation

ℹ *Full online documentation here*

Due to experimental constraints, the number of separated color channels that can be acquired in the same imaging batch is limited. In some experiments, biologists prefer to stain and acquire both the nucleus and the cell junctions in the same color channel. But the segmentation of the biological structures needs the two signals to be separated.

##### 2.8.1 Morphological separation

In FishFeats, we proposed a first option to separate these structures based on their morphological differences (big round volumes for nuclei and linear structures for junctions). By applying morphological filtering to enhance linear or spherical structures, the two signals can be separated in two virtual channel images.

##### 2.8.2 `SepaNet` : Deep learning separation

Separation by morphological filtering necessitated to tune parameters to adapt them to new image resolutions or biological structure sizes. Thus, we introduced a new neural network,

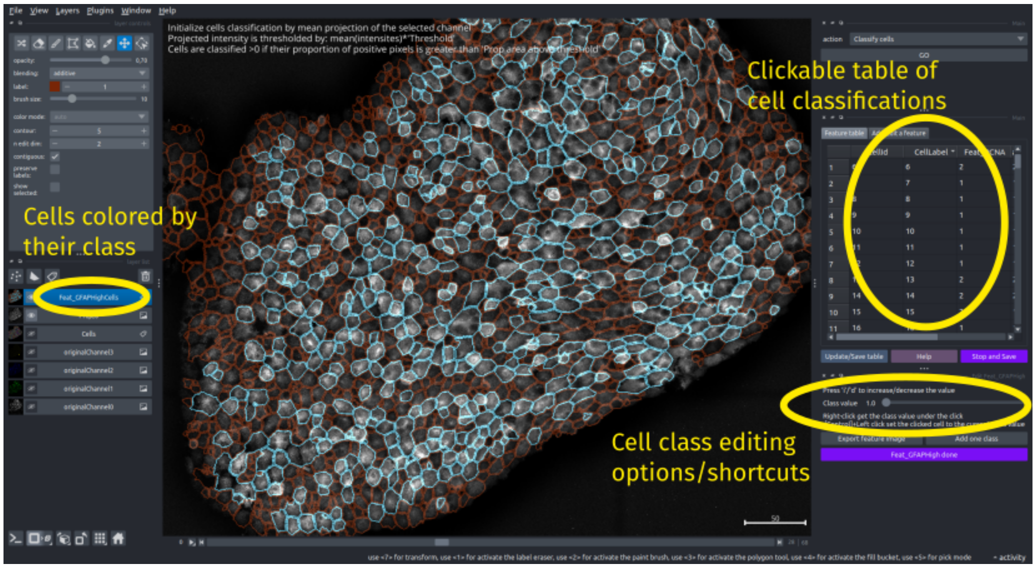

**Fig 11.** *Interface to edit cell classification for a user-defined feature (e.g. PCNA positive).*

`SepaNet` , that we implemented and trained to separate the two biological structures. `SepaNet` follows a U-Net [Ronneberger *et al.*, 2015] architecture, but after the most condensed layer part, two upsampling paths are done in parallel, one for each output image (Fig 12).

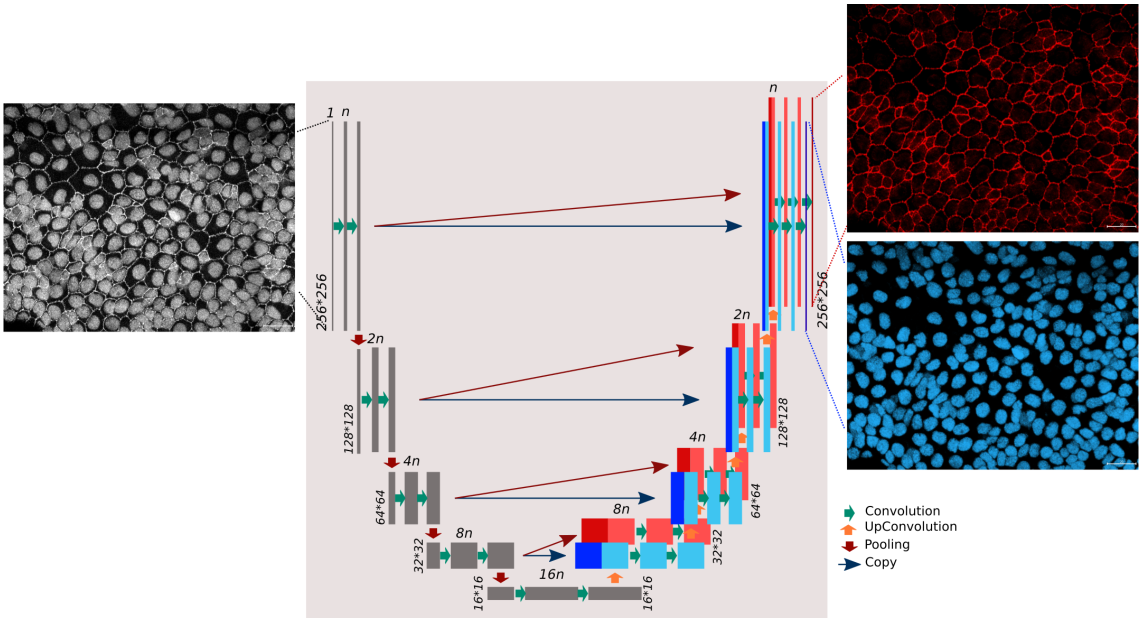

**Fig 12.** `SepaNet` *architecture, derived from U-Net architecture with two parallel upsampling parts.*

It was implemented with `tensorflow`, and the available network (on our repository) is compatible with `Keras version 2` (not with the version 3). In future developments, we intend to reimplement it in `Pytorch` for more compability.

##### 2.8.3 SepaNet training dataset

For the training dataset, we had some images containing either the junctional or nuclear staining in one channel, and one other channel with both stainings combined, which were the optimal conditions. We had more images that contained only the two signals in two separated channels. We took advantage of these additionnal data by normalizing and mixing these separated images to create a virtual mixed channel. The intensity normalization to combine the two channels together was varied to create more data as a form of data augmentation.

##### 2.8.4 SepaNet in FishFeats

In *FishFeats* , `SepaNet` can be used through the `separate junction and nuclei` steps. The trained model is available in our repository and can be directly downloaded from the github repository. `SepaNet` can easily be retrained to fit new imaging conditions or separate other biological structures with a new training dataset, outside of *FishFeats* (no option is proposed yet for it).

After the separation has been caclulated (Fig 13), *FishFeats* will directly use the virtual channel for the other steps (cell or nuclei segmentation). Note however that intensity measurement should not be done on the separated staining, as the exact pixel value of the virtual channels are generated by the neural network.

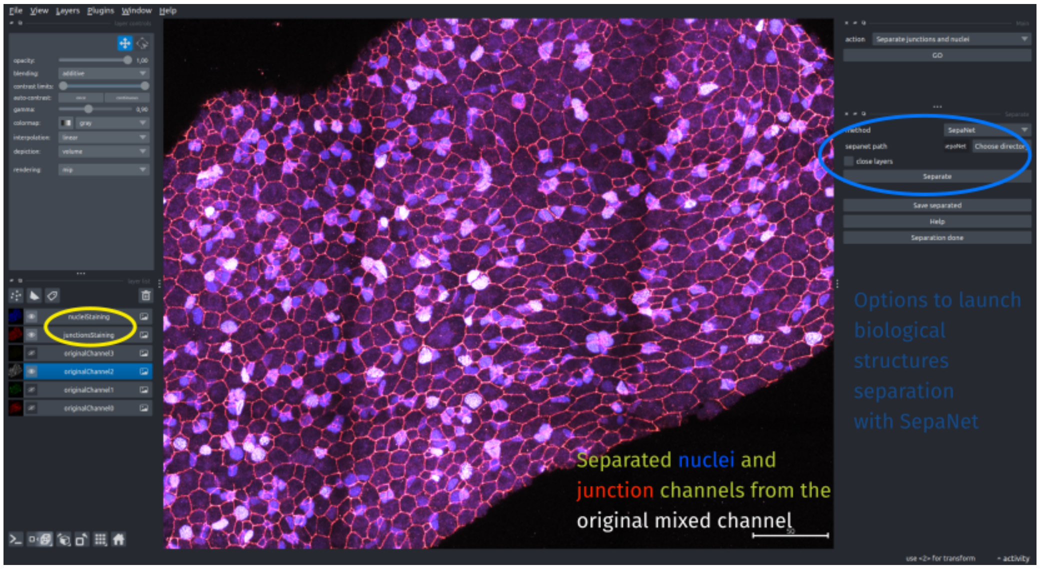

**Fig 13.** *FishFeats option to separate biological structures acquired in the same color channel.* `SepaNet` *creates two virtual color channels, containing the nuclei (in blue) and the junctions (in red) from the original mixed channel (in white).*

#### 2.9 Other options

During FishFeats development, we added several basic options based on users feedback/wishes. Notably:

- An option to perform hierarchical clustering of the cells, based on the different measurements (cytoplasmic, RNA counts...) and displaying the clustered cells directly in the plugin
- Making the cell segmentation compatible with napari-griottes plugin

- Adding a background grid to the viewer to have a landmark in the visualization
- Preprocessing the nuclear staining to improve segmentation performance (with the possibility to use noise2void denoising [Krull *et al.*, 2019])

Please refer to the plugin interface and the online documentation at https://gletort.github.io/FishFeats/ for more information.

### 3 Results table

All measured features (cell size, position, nucleus size, position, cytoplasmic/nuclear intensity, RNA counts (DNA/RNA/Protein dots), cell classification..) are grouped in a single *.csv* table, where each line contains the joint information of one cell (Figure 14. This table is called *imagename_results.csv* and is saved in the 'results' directory. It is ready to be used in any software for table quantification as Excel, Prism, R, Python for downstream analysis.

| | B | C | D | E | F | G | H | I | J | K | L | M | N | O | P | Q | R | S | T |
|---|---|---|---|---|---|---|---|---|---|---|---|---|---|---|---|---|---|---|---|
| 1 | CellLabel | XPosPi | YPos | ZPosPi | XPosMicro | YPosMi | ZPosMi | CellAreaM | Nucle | NucleusLa | NucleusVolume | NucleusZPos | NucleusZ | Nucle | Nucleus | YPos | nbRNA_C2 | nbRNA_C1 | nbRNA_C3_Load |
| 2 | 18 | 1429.6 | 43.63 | 13 | 295.925154 | 9.03236 | -49999.5 | 147.40056 | 18 | 18 | 255.465738 | 2.700645756 | 5.401292 | 1429 | 47.4237 | 0 | 2 | 1 | 6 |
| 3 | 21 | 875.66 | 37.08 | 15 | 181.261327 | 7.67548 | -49999.5 | 130.77515 | 21 | 21 | 218.701296 | 3.375195925 | 6.750392 | 854.6 | 14.6419 | 0 | 22 | 4 | 4 |
| 4 | 24 | 1337.4 | 28 | 14 | 276.844021 | 5.80327 | -49999.5 | 43.920225 | -9999 | -9999 | -9999 | -9999 | -9999 | -9999 | -9999 | 0 | 1 | 0 | 3 |
| 5 | 25 | 1382.6 | 39.66 | 14 | 286.191416 | 8.20965 | -49999.5 | 91.782558 | 25 | 25 | 220.6937745 | 4.381128046 | 8.762256 | 1383 | 40.9208 | 0 | 0 | 0 | 4 |
| 6 | 27 | 1210.6 | 49.84 | 14 | 250.593986 | 10.3169 | 7 | 239.8687 | 27 | 27 | 296.600778 | 5.888868824 | 11.77774 | 1210 | 47.6134 | 0 | 2 | 6 | 3 |
| 7 | 28 | 1002.6 | 51.08 | 15 | 207.536774 | 10.573 | -49999.5 | 140.58757 | 28 | 28 | 243.4037445 | 5.71138104 | 11.42276 | 1004 | 50.7542 | 0 | 2 | 3 | 11 |
| 8 | 29 | 765.09 | 33.49 | 2 | 158.373861 | 6.93332 | -49999.5 | 18.810711 | 29 | 29 | 100.5880275 | 1.375505857 | 2.751012 | 771.5 | 27.7184 | 0 | 0 | 0 | 1 |
| 9 | 30 | 1118.7 | 61.68 | 15 | 231.562317 | 12.7675 | -49999.5 | 258.25092 | 30 | 30 | 266.1137145 | 7.060220594 | 14.12044 | 1132 | 65.2397 | 0 | 2 | 5 | 7 |
| 10 | 31 | 1465.9 | 54.53 | 13 | 303.441089 | 11.2871 | -49999.5 | 54.718173 | 31 | 31 | 129.6824985 | 1.0337849 | 2.06757 | 1464 | 56.1145 | 0 | 0 | 0 | 0 |
| 11 | 32 | 964.64 | 41.48 | 15 | 199.679919 | 8.58664 | -49999.5 | 12.640455 | 32 | 32 | 151.0641495 | 5.704297263 | 11.40859 | 960.4 | 7.93845 | 0 | 1 | 0 | 2 |
| 12 | 33 | 1279.5 | 63.9 | 14 | 264.85323 | 13.2272 | -49999.5 | 108.49367 | 33 | 33 | 323.2742805 | 5.674497979 | 11.349 | 1277 | 60.0714 | 0 | 1 | 0 | 1 |
| 13 | 34 | 1039.5 | 43.34 | 15 | 215.167091 | 8.97063 | -49999.5 | 4.71339 | 34 | 34 | 112.4572005 | 1.436559345 | 2.873119 | 1046 | 53.2764 | 0 | 3 | 0 | 3 |
| 14 | 35 | 1054.4 | 41.5 | 15 | 218.26731 | 8.5905 | -49999.5 | 10.626552 | 35 | 35 | 126.361701 | 5.281536114 | 10.56307 | 1077 | 9.93896 | 0 | 0 | 1 | 4 |
| 15 | 37 | 1339.7 | 74.35 | 14 | 277.311882 | 15.39 | -49999.5 | 177.60911 | 37 | 37 | 290.3233995 | 5.892332669 | 11.78467 | 1333 | 72.2872 | 0 | 2 | 6 | 2 |
| 16 | 39 | 954.67 | 59.87 | 15 | 197.616524 | 12.394 | -49999.5 | 39.506778 | 39 | 39 | 169.3606725 | 3.584313725 | 7.168627 | 974.1 | 40.0798 | 0 | 1 | 0 | 3 |
| 17 | 40 | 690.73 | 66.14 | 2 | 142.982064 | 13.6901 | -49999.5 | 51.504498 | 40 | 40 | 138.9807315 | 0.986203176 | 1.972406 | 696 | 65.1714 | 0 | 0 | 0 | 0 |
| 18 | 41 | 735.39 | 69.38 | 2 | 152.225193 | 14.3615 | 1 | 97.995663 | 41 | 41 | 246.081807 | 2.645307331 | 5.290615 | 736.9 | 69.2473 | 0 | 0 | 0 | 0 |
| 19 | 42 | 915.78 | 75.77 | 15 | 189.567366 | 15.6853 | -49999.5 | 109.69344 | -9999 | -9999 | -9999 | -9999 | -9999 | -9999 | -9999 | 0 | 2 | 3 | 0 |
| 20 | 43 | 1048.9 | 62.82 | 15 | 217.113955 | 13.0032 | 7.5 | 43.363188 | 43 | 43 | 493.534782 | 4.306976038 | 8.613952 | 1059 | 52.8826 | 0 | 10 | 0 | 11 |
| 21 | 44 | 1497 | 72.78 | 13 | 309.874372 | 15.0646 | -49999.5 | 78.585066 | 44 | 44 | 123.4265445 | 0.907568131 | 1.815136 | 1499 | 73.1403 | 0 | 0 | 0 | 3 |

**Fig 14.** *FishFeats results table containing all the measures done and joint together in a cell-by-cell table.*

### 4 Experimental data

The experimental data presented in the corresponding Application Note and this supplementary data correspond to 3D confocal images of dorsal views of dissected telencephala from 4 months post-fertilization (mpf) zebrafish. After fixation the brain was stained by whole-mount immunohistochemistry (IHC) for Zo1 (Zonula occludens 1) and Sox2 revealed using a unique secondary antibody coupled to the fluorophore Alexa 405 [Ortica *et al.*, 2025, Mancini *et al.*, 2023]. Expression of *pcna, her4* and *hey1* (in green, orange and red) were detected using for fluorescent in situ hybridization (FISH) (RNAscope from BioTechne). Images were acquired using a LSM980 confocal microscope (Zeiss) using a 40X oil objective (Plan-Apochromat 40×/1.3 Oil M27). Bit depth for all the images acquired was 16 bit, a tile scan of multiple Z stacks all with a voxel size of 0.207 µm by 0.207 µm by 0.5 µm. Zo1 outlines the apical cell contours, while Sox2 is used for the identification of neural stem cells and progenitor cells corresponding mostly to the first layer of cells.
